# Interactions between specific breeding system and ploidy play a critical role in increasing niche adaptability in a global food crop

**DOI:** 10.1101/2020.09.09.290429

**Authors:** Nathan Fumia, Daniel Rubinoff, Rosana Zenil-Ferguson, Colin K. Khoury, Samuel Pironon, Michael A. Gore, Michael B. Kantar

## Abstract

Understanding the factors driving ecological and evolutionary interactions of economically important plant species is important for sustainability. Niches of crop wild relatives, including wild potatoes (*Solanum* section *Petota*), have received attention, however, such information has not been analyzed in combination with phylogenetic histories, genomic composition and reproductive systems. We used a combination of ordinary least-squares (OLS) and phylogenetic generalized least-squares (PGLM) analyses to identify the discrete climate classes that wild potato species inhabit in the context of breeding system and ploidy. Self-incompatible diploid or self-compatible polyploid species significantly increase the number of discrete climate niches inhabited. This result was sustained when correcting for phylogenetic non-independence in the linear model. Our results support the idea that specific breeding system and ploidy combinations increase niche divergence through the decoupling of geographical range and niche diversity, and therefore, these species may be of particular interest for crop adaptation to a changing climate.

## Introduction

Potato (*Solanum tuberosum* L.) is the most important tuber crop worldwide and is the fourth most important crop internationally (Castañeda-Álvarez et al. 2015). However, there is a lack of genetic diversity among many crops, including *S. tuberosum* (Jansky et al., 2013; Khoury et al., 2014), placing increased pressure upon crop management protocols and food security. A proven approach to increasing genetic diversity in crop species is through the utilization of wild relatives for crop improvement (Jansky et al., 2013; Mehrabi et al., 2019). Cultivated potato has 199 known wild relatives, forming the *Solanum* section *Petota*, inhabiting 16 countries in the Americas, and ranging from 38° N to 41° S (Hijmans, 2001); 72 of the most threatened and useful species to humans have recently been prioritized for conservation (Castañeda-Álvarez et al., 2015). These 72 species are most commonly found in tropical highlands at 600 to 1200 m in elevation and possess phenotypes similar to cultivated potato through the production of a starchy tuber (Hijmans, 2002).

Given the importance of maintaining the crop’s productivity, many attributes of the wild relatives of *S. tuberosum* have been defined, including their ploidy, breeding system, germplasm classification, endosperm balance number, single and multi-gene phylogenies, and geographic ranges (Hijmans, 2001; Spooner, 2001; Spooner, 2007; Castañeda-Álvarez et al., 2015; Robertson et al., 2011). These data can be used to discover novel beneficial characteristics present within the wild relative germplasm such as biotic and abiotic resistances as well as to quantify trait introgression. Furthermore, research has identified potato as one of the crops in Sub-Saharan Africa with the highest potential to benefit from crop wild relatives for climate change adaptation, however, these results have not been integrated with biological (e.g. breeding system and ploidy) and evolutionary (e.g. phylogenetic tree) information (Pironon et al., 2019). Despite the wide array of information surrounding the wild relatives of potato, one attribute continues to be under-defined - the discrete climate zones (e.g. niche) each species inhabits and the factors involved (e.g., breeding system, ploidy) in driving the evolution of the highly dynamic climatic diversity in *Solanum* section *Petota*.

Individually exploring life history traits (Wendel and Cronn, 2003; Hijmans et al., 2007; Köhler et al., 2010; Sessa, 2019) such as the breeding system has led to contradictory conclusions regarding these traits’ influence on ecological niche (Peterson et al., 1999; Husband et al., 2008; Robertson et al., 2010; Campbell, 2013; Grossenbacher et al., 2016; Park et al., 2017; Grant, 2020), while exploring other traits such as ploidy (Baniaga et al., 2020) has shown a consistent influence. For example, diversification models Zenil-Ferguson et al. (2019) showed that ploidy is the most probable pathway to evolve self-compatibility across Solanaceae. Therefore, there exists an important interaction between ploidy and breeding system (Barringer, 2007; Husband et al., 2008; Robertson et al., 2010) that might impact evolutionary and ecological processes (Sessa, 2019). Furthermore, polyploidization facilitates self-compatibility because whole genome duplication provides security against inbreeding depression (Barringer, 2007; Husband et al., 2008; Robertson et al., 2010; Zenil-Ferguson et al., 2019); whereas, self-compatible diploid populations often suffer from large inbreeding depression (Barringer and Geber, 2008; Husband and Schemske, 2017). As a result, diploid populations are more reliant on self-incompatibility to drive adaptive changes. In *Solanaceae*, polyploid species show higher rates of self-compatibility (Barringer, 2007; Husband et al., 2008; Robertson et al., 2010). This clear interaction between ploidy and breeding systems provides the opportunity to test two key hypotheses: first, that self-compatible species rely on polyploidy in order to generate the variation they need to colonize diverse ecological niche space; and second, that diploid species rely on outcrossing to increase niche breadth through gene flow.

To identify the driving factors of ecological diversity in potato wild relatives, we investigated two biological aspects of ecological diversity - breeding system and ploidy in 72 wild relatives of potato. We combined species’ occurrence, climatic, biological (e.g. breeding system and ploidy), and phylogenetic tree of *Solanum* taxa to test whether the niche diversity of a given species is guided by a specific breeding system and ploidy interaction. To account for the potential decoupling of geographical range and niche breadth (Randel et al., 2009), the measure of climatic diversity is through the use of discrete climate-classification of each occurrence of these wild relative species. This work supports classic ecological theory of niche divergence without the requirement of inferring continuous species distributions from point-based climate descriptions by featuring the relationship between two common intrinsic factors of niche expansion: (1) decreased reliance on outcrossing reproduction of polyploid variants; and (2) increased reliance on outcrossing reproduction of diploid variants (Roughgarden, 1972; Barton, 1996; De Bodt et al., 2005; Johnson et al., 2014).

## Materials and Methods

### Data Collection

Data organization and analyses were conducted using the R (R Core Team, 2020) packages “raster” (Hijmans, 2020) and “tidyverse” (Wickham et al., 2019). We obtained 49,165 occurrence records of the 72 *Solanum* species sourced from Castañeda-Álvarez et al (2015). These occurrences represent the most threatened and useful wild relatives of *Solanum tuberosum*, the previously cleaned points were further filtered for those lacking latitudinal and/or longitudinal information, resulting in a total of 37,032 total occurrence points (Castañeda-Álvarez et al., 2015). Next, the Köppen-Geiger three-tier climate class system was acquired from Rubel and Kottek (2010). The Köppen-Geiger climate class system divides climates into five main groups that are subdivided based on seasonal precipitation and temperature that result in 30 potential discrete classes globally (reviewed in Rubel and Kottek, 2010). Three-tier climate classes were extracted at each occurrence point. The total number of climate classes per species was counted for each species and climate classes with three or fewer occurrences were removed in order to avoid “by-chance” occurrences. Using discrete climate classes allows for a single measure of both niche diversity and breadth. See github repository “https://github.com/Nfumia/Potato_nichediversity_drivers“for code and data files.

### Linear Models for Climate Classes

A linear model was fitted using R package “stats” (R Core Team, 2020) to identify which interaction of biological factors is correlated with niche diversity in *Solanum* section *Petota*. We used the number of discrete climate classes in which each taxon can occur as a proxy for niche breadth, as these niches vary spatially within the five broad descriptors of tropical, dry, temperate, continental, and polar each of which possessing 2-12 subclassifications. For example, *S. stoloniferum* has fifteen discrete niches in which it occurs, but *S. albornozii* has only one, a temperate oceanic environment. The number of discrete climate classes is the response variable for the model. The predictor variables were combinations of ploidy (Castañeda-Álvarez et al., 2015) and breeding system (Robertson et al., 2010; Zenil-Ferguson et al., 2019) for each species, which were coded as dummy variable interaction terms: self-incompatible diploid, self-compatible diploid, self-compatible asexually propagating polyploid, and unknown breeding system asexually propagating diploid.

### Phylogenetic Tree and Phylogenetic Linear Models

A Bayesian molecular clock phylogeny with time-calibration of section *Petota* to outgroups of domesticated tomato (*Solanum lycopersicum*) and domesticated eggplant (*Solanum melongena*) was estimated using 32 plastid genomes and compared to the most recent time-calibrated phylogeny of Särkinen et al. (2013). Due to a lack of plastid genome availability for some species in *Solanum* section *Petota*, only 27 of the 72 prioritized wild relative species were present in our subsequent analyses. Furthermore, 32 species (27 potato wild relatives, 2 domesticated potato, 1 domesticated tomato, 1 tomato wild relative, 1 domesticated eggplant) were aligned using the software MAFFT (multiple alignment using fast Fourier transform) via maxiterate version (Katoh, 2009). MrBayes (Huelsenbeck and Ronquist, 2001) as implemented in the Geneious software package (Kearse et al., 2012) was used to conduct an initial phylogenetic analysis (Vallejo-Marín and O’Brien, 2006; Newton et al., 1999). We used a chain length of 10 million generations with 25% (or 2.5 million) burn in and a subsampling frequency every 1,000 generations. The General Time Reversible (GTR) substitution model was employed for the Bayesian analysis with rate variation of gamma, including 4 categories.

We used the Bayesian uncorrelated relaxed clock-model dating method as implemented in BEAST2 (Bouckaert et al., 2019). The uncorrelated relaxed clock-model allows for rate variation across branches and measures for rate autocorrelation between lineages. Node ages are estimated simultaneously in BEAST2, and, therefore, uncertainty is incorporated into the node-age estimation. Our Bayesian MCMC tree output was used as a starting phylogeny. The Hasegawa-Kishino-Yano (HKY) model for DNA base pair substitution was used to better estimate the substitution rates of transition versus transversion as well as the Felsenstein (F81) proposed four-parameter model. A Kappa of 2.0, as estimated by BEAUti2 (Bouckaert et al., 2019), was employed. Calibration points for the node-age estimation were sampled from Särkinen et al. (2013) to create calibration priors: (1) tomato – potato split circa 8 mya (95% HPD 7—10), and (2) eggplant – tomato/potato split circa 14.3 mya (95% HPD 13-16). These calibration points reflect a normal distribution with standard deviations of 0.85 and 1.10 million years, respectively. Yule tree prior with uniform distribution was used given all ingroup and outgroup species in this study currently persist *ex-situ* and/or *in-situ*. Priors were manually generated for each monophyletic clade showing greater than 85% posterior probability from the MrBayes MCMC analysis. Default priors were used for all other parameters. A total of 100 million generations, 10 runs with 10 million generations each, were run in BEAST2 (Bouckaert et al., 2019).

Using the time-calibrated phylogeny (Supplemental Fig. 1), we estimated the phylogenetic generalized linear models version of the OLS models proposed in the previous section to account for potential phylogenetic signals in the errors (Felsenstein, 1985; Hansen, 1997). This is an important step, since it is possible that our explanatory variables are not tracking the evolutionary history of the *Petota* section, and can incorrectly conclude strong correlations between the climatic classes and the life history traits (Uyeda et al., 2018).

These phylogenetic linear models were estimated using a maximum likelihood PGLM with the R package “phylolm” (Ho and Ane, 2014). For all the PGLMs we assumed a Brownian motion model of evolution (Grafen, 1989; Martins and Hansen, 1997; Revell and Harmon, 2008). Outgroup species and cultivated potato were removed at this point due to the inability to differentiate between cultivated and wild occurrence of the given species. This resulted in retention of 27 potato wild relative species, comprising the four major monophyletic clades of section *Petota* (Spooner et al., 2014), for use in the PGLMs analysis.

## Results

### Climate Regression

The 72 prioritized species in the *Solanum* section *Petota* examined here occurred in 17 distinct climates with individual species distributions ranging from a single climate (e.g. *S. albornozii, S. chilliasense, S. lesteri*) to 15 distinct climates (e.g. *S. stoloniferum*). Within this range exists a spectrum of breeding system and ploidy combinations between and within these species and their populations, exhibiting different extents of climate niche diversity (Fig. 1). This analysis showed that distinct breeding system and ploidy combinations existed in a different number of niches (p = 3.4 × 10^−7^), described as the number of discrete Köppen-Geiger climate classes. Species that possess populations that are self-incompatible diploid and self-compatible polyploid show the greatest mean climate diversity with 11 discrete climate classes (Fig. 1). Self-incompatible diploid species exhibit a greater average niche diversity when compared to self-compatible diploid species (Fig. 1). Furthermore, diploid species possessing populations showing polyploidization demonstrate greater sustained ecological divergence.

**Figure 1.**
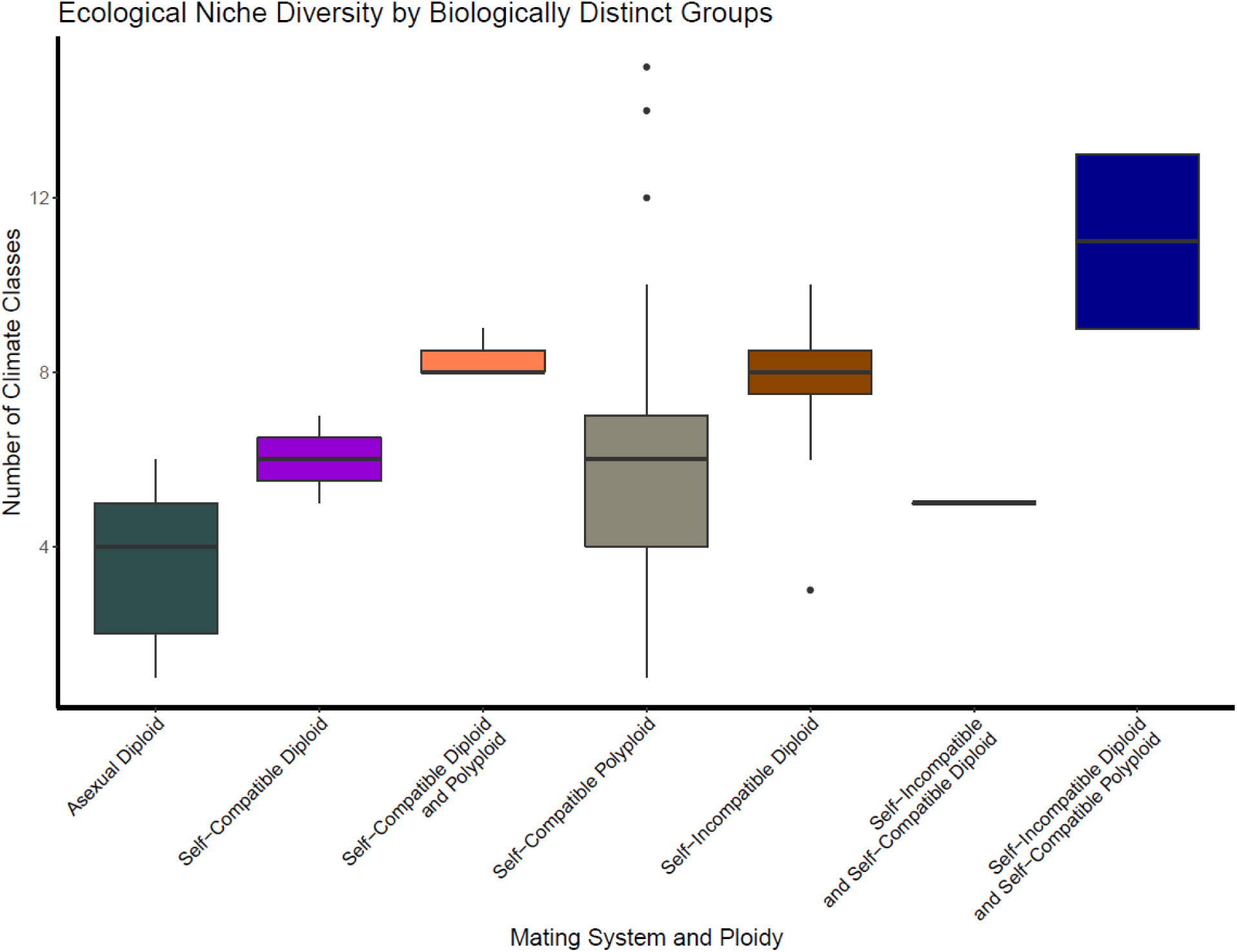
Boxplot of niche diversity by breeding system and ploidy interaction in potato wild relative species. Many species exist containing multiple subpopulations with differing biological factors, as seen by combination of such factors on the x-axis.

The maximum likelihood intercept value of ecological niche diversity is 2.81±1.01 climate classes. Species existing as self-incompatible diploid or self-compatible polyploid have significantly (p-value < 0.01) larger climatic niches by 3.13±0.73 and 3.62±0.79 discrete climate classes, respectively (Table 1). However, other predictor (self-compatible diploid, asexually propagating unknown breeding system diploid) variable slope values are not significantly different from zero, and, therefore, they exert no measurable influence on niche diversity within *Solanum* section *Petota*. Overall, the model explained a moderate amount of variance with an adjusted R-square of 0.39.

**Table 1.** Results from the linear model for climatic niche diversity following Gaussian distribution. The number of discrete climate classes in which each taxon can occur (i.e. a proxy for niche breadth) is the response variable, *Climate Niche Diversity*. The predictor variables are combinations of ploidy and breeding system for each species, which were coded as dummy variable interaction terms: self-incompatible diploid, self-compatible diploid, self-compatible asexually propagating polyploid, and unknown breeding system asexually propagating diploid.

| **Ordinary Least-Squares Results** |  |
| --- | --- |
|  | *Dependent variable:* |
|  | Climatic Niche Diversity |
| Self-Incompatible Diploid | 3.134***<br>(0.734) |
| Self-Compatible Diploid | 0.569<br>(1.021) |
| Self-Compatible Polyploid (Asexual) | 3.624***<br>(0.789) |
| Asexual Diploid | 0.883<br>(0.992) |
| Constant | 2.813***<br>(1.006) |
| Observations | 72 |
| R<sup>2</sup> | 0.426 |
| Adjusted R<sup>2</sup> | 0.392 |
| Residual Std. Error | 2.493 (df = 67) |
| F Statistic | 12.424*** (df = 4; 67) |
| Note: | * ** ***<br>p p p<0.01 |

### Evolutionary Climate Regression

In the PGLMs fitted using our estimated time-calibrated phylogeny (Fig. 2), we found an estimated intercept value of 6.43±1.67 (Table 2). The PGLMs confirmed the correlations of OLS models, with self-incompatible diploid (3.98±1.04) and self-compatible polyploid (2.57±0.98) significantly increasing ecological diversity (Table 2). As with OLS, the other predictor variables in PGLMs are not significantly different from zero.

**Table 2.** Results from phylogenetic linear models for climatic niche diversity following Brownian motion. The number of discrete climate classes in which each taxon can occur (i.e. a proxy for niche breadth) is the response variable, *Climate Niche Diversity*. The predictor variables are combinations of ploidy and breeding system for each species, which were coded as dummy variable interaction terms: self-incompatible diploid, self-compatible diploid, self-compatible asexually propagating polyploid, and unknown breeding system asexually propagating diploid.

| Phylogenetic Least-Squares Results |  |
| --- | --- |
|  | *Dependent variable:* |
|  | Climatic Niche Diversity |
| Self-Incompatible Diploid | 3.984***<br>(1.036) |
| Self-Compatible Diploid | 1.856<br>(1.536) |
| Self-Compatible Polyploid (Asexual) | 2.574*<br>(0.975) |
| Asexual Diploid | -2.332<br>(1.620) |
| Constant | 6.426***<br>(1.670) |
| Sigma<sup>2</sup> | 8.088e-09<br>(2.967e-09,1.067e-08) |
| Sigma<sup>2</sup> Error | 4.497<br>(1.650,5.933) |
| Observations | 27 |
| R<sup>2</sup> | 0.527 |
| Adjusted R<sup>2</sup> | 0.441 |
| Residual Std. Error | 4.761 (df = 22) |
| Parametric Bootstraps | 100 |
| *Note:* | * ** ***<br>p p p<0.01 |

**Figure 2.**
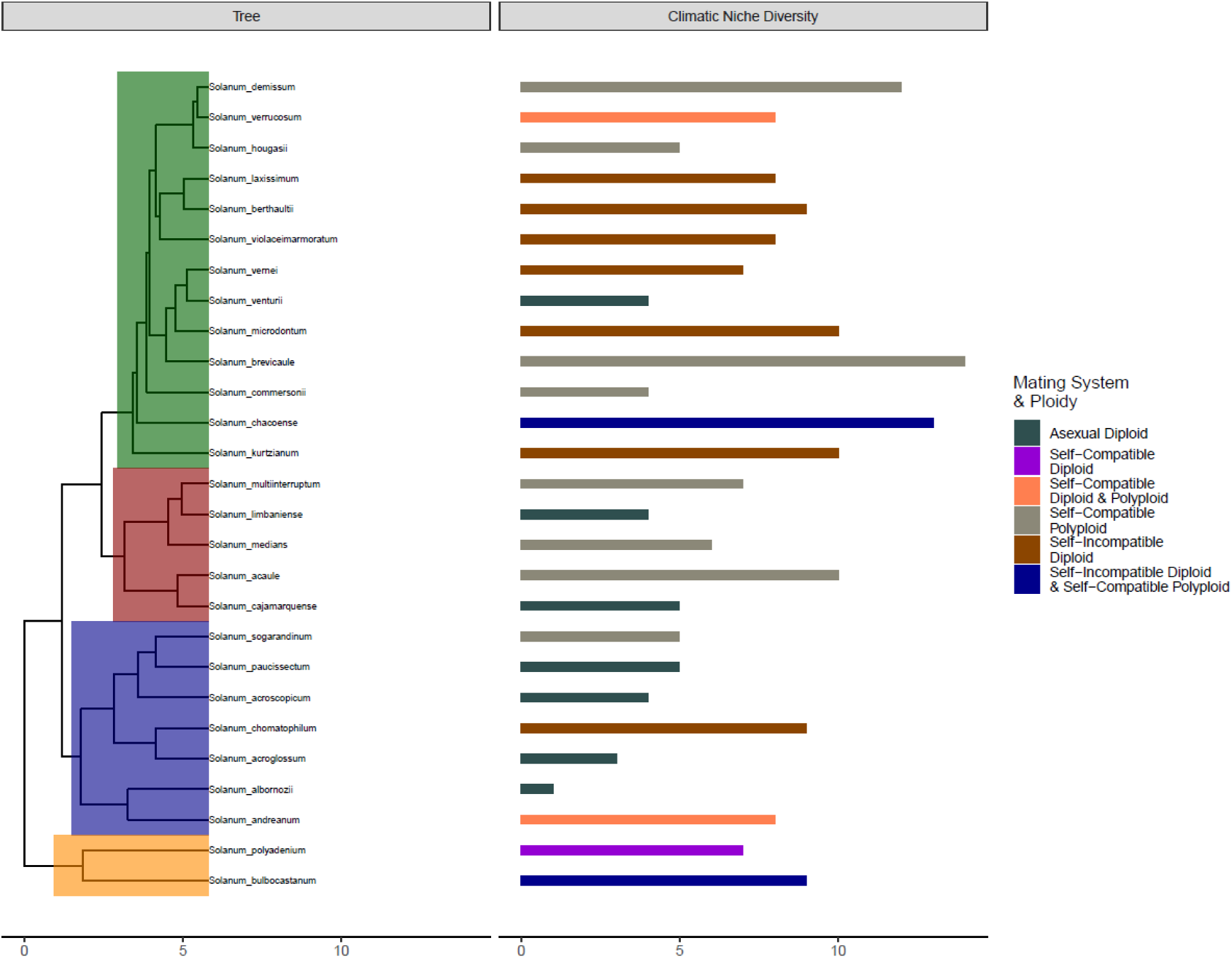
Dual figure with time-calibrated molecular clock phylogeny (left) with climatic niche diversity (i.e. number of climate classes occupied) (right). On the left side, the x-axis scale bars represent millions of years and the background coloration of the phylogenetic tree highlights widely accepted clades of *Solanum* section *Petota*. On the right side, the number of climate classes a species occurs in is represented by the size of the horizontal bar and measured with the x-axis scale bar and the coloration of the horizontal bars represent species biological attributes as breeding system with ploidy.

## Discussion

Clarifying the impacts of plant traits on niche divergence is important to understanding the structure of global patterns of biodiversity and evolution in plant lineages (Cavender‐Bares et al., 2009). Furthermore, life history traits can provide clues about the potential resiliency of plants with increased development of wild areas leading to changes in habitat and climate for many species. However, resilience may be tightly linked with other characteristics. In *Solanum* section *Petota*, the interaction of two specific characters, breeding system and ploidy, explain a large portion of the variation in niche divergence. The models presented here, OLS and PGLMs, explain 39% and 44% (R-squared), respectively, of the climatic niche variation present within *Solanum* section *Petota* with two alternate ends of the biological spectrum serving as the most significant predictors. On one end, self-incompatible diploid species exhibit the greatest significant correlation to climatic niche diversity within potato wild relatives. Such sustained diversity is likely the result of constant capacity for outcrossing between these species and their subsequent heterogenous design, fashioning an adaptive and resilient population through long-distance gene flow (Loveless and Hamrick, 1984). Due to the interaction between ploidy and breeding system, self-incompatible diploid species show niche diversity similar to self-compatible polyploid species, confirming the dynamic nature of the *Solanaceae* system (Barringer, 2007; Husband et al., 2008; Robertson et al., 2010; Zenil-Ferguson et al., 2019). However, self-fertilizing polyploid species have a short-term advantage as they can colonize new environments with very few individuals.

For all the Solanaceae family self-incompatible diploid has been shown to be the ancestral state (Zenil-Ferguson et al., 2019), they also have faster net diversification compared to all self-compatibles, both diploid and polyploid (Wright et al., 2013). The expectation given the success of these lineages in diversification is that self-incompatible diploids should have broader niches, an unexpected result was that self-compatible polyploids diversified in a similar way. Evolutionarily this may be a temporal effect, polyploids are successful in short time scales and this may explain the success in diversification identified here, however, this study does not disentangle evolutionary time-scales. Our results suggest that self-compatible diploids appear evolutionarily transient, and the evolution of self-compatibility appears to occur very rarely without a polyploidy event in Solanaceae. This suggests that polyploidy is just an evolutionary byproduct of trying to become self-compatible, allowing for rapid establishment in many new environments.

Self-compatible polyploid species have increased climatic niche diversity which, given their increased genetic variation and plasticity through additional sets of chromosomes, make them capable of adaptive and resilient population generation (Soltis and Soltis, 1999). Polyploidy allows self-fertilizing section *Petota* species to maintain and derive novel diversity typically observed in outcrossing/self-incompatible diploid populations. These differences between breeding system and ploidy with niche diversity provide support for the use of these variable combinations as driving evolutionary forces, with qualitative results (Fig. 1) being supported by OLS (Table 1) and PGLMs (Table 2).

Our results suggest the potential to use ecologically plastic species to enhance the adaptability of cultivated potato lines in the face of climate change. The ultimate goal of this investigation is increased beneficial genetic variation among cultivated potato varieties developed through introgression of the various wild adaptations. However, the wild species have limited cross-compatibility with *S. tuberosum*, as evidenced in their endosperm balance numbers. Therefore, time is needed in order to operationalize this diversity in agricultural fields, so that favorable environmental adaptations from a subset of ecologically plastic species, can be introgressed while breaking linkages to agronomically unfavorable traits.

The impact of breeding system on the evolution of climatic niche diversity amongst plants is still unclear and the *Solanum* section *Petota* system contributes important evidence for a multilayered role where breeding system and ploidy interact synergistically with one another. In one case, self-incompatible breeding systems play a large role in sustaining niche diversity over time (Park et al., 2017) when species are diploid, possessing limited reproductive barriers. In contrast, self-compatible breeding systems comparatively increase niche diversity when species are polyploid, by enhancing their ability to reach, reproduce, establish, and adapt (Campbell, 2013) with the biological safety net of increased “buffering capacity” through genetic variation (Wendel and Cronn, 2003). Further investigations could focus on the decoupling of breeding system and ploidy; however, due to the self-incompatibility conferred by S-RNases found in polyploid populations of *Solanaceae* this is challenging (Robertson, 2010; Barringer, 2007; Husband et al., 2008). Furthermore, this study was not able to completely decouple ploidy and breeding system interactions due to lack of data on particular species’ breeding systems, exemplifying the need for more than DNA collection. Additionally, a limitation of this analysis is the limited number of species available for PGLMs, which was due to a lack of publicly available plastid genome sequence data.

Increasing effective genetic diversity through polyploidization has the potential to increase the number of niches to a similar extent as would occur with an outcrossing diploid population. The breeding system is the main driver of niche divergence in the self-incompatible diploid populations, while only a secondary contributor in the self-compatible polyploid populations. Despite the biological differences, the resulting niche diversity is not seen in a difference of preferred climate type but rather the extent of climatic diversity (Supplemental Fig. 2). Through decoupling geographical range size and niche breadth (Randel et al., 2009), this study tests classic theory by utilizing a highly diverse, economically important section of plants. Our findings lend credence to the hypothesis that these ecologically plastic responses evolved over millions of years in species with populations of self-incompatible diploids and self-compatible polyploids, and, therefore, these species should be prioritized for conservation and for use to adapt our cultivated varieties to a changing climate.

## Acknowledgements

We would like to the Hawaii Agriculture Research Center and the University of Hawaii Office of Sustainability for their support of Nathan Fumia through the Sustainable Agriculture Fellowship, the Information Technology Systems at the University of Hawai’i at Manoa for computer processing support, and access to data via the Centro Internacional de la Papa. We would like to thank Cornell University for supporting the sabbatical of Dr. Michael A. Gore to contribute to this manuscript.

## Supplemental Figures

**Supplemental Figure 1.**
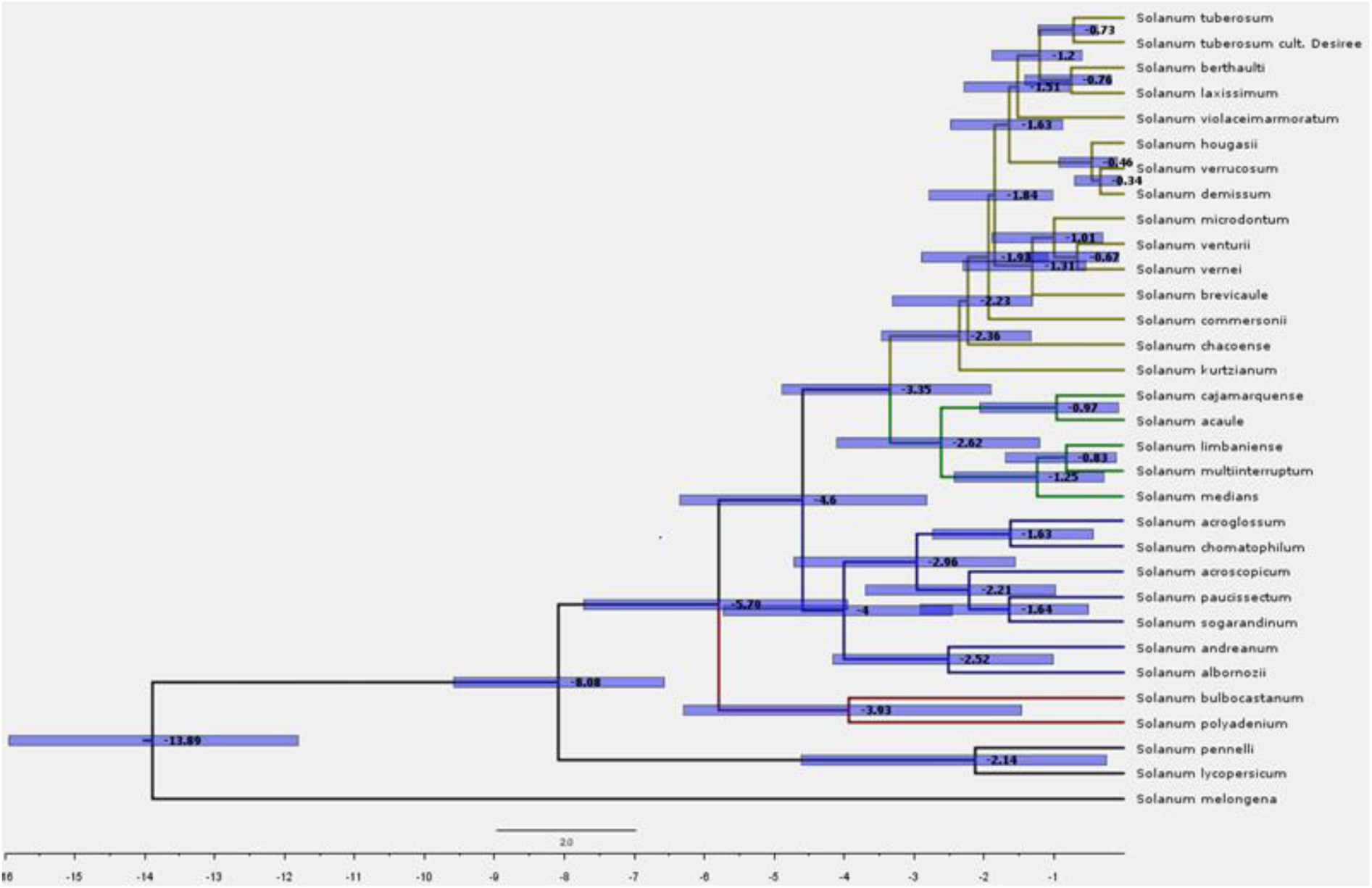
Time-calibrated uncorrelated relaxed molecular clock phylogeny with outgroups and cultivated varieties of potato included as generated with BEAST2.

**Supplemental Figure 2.**
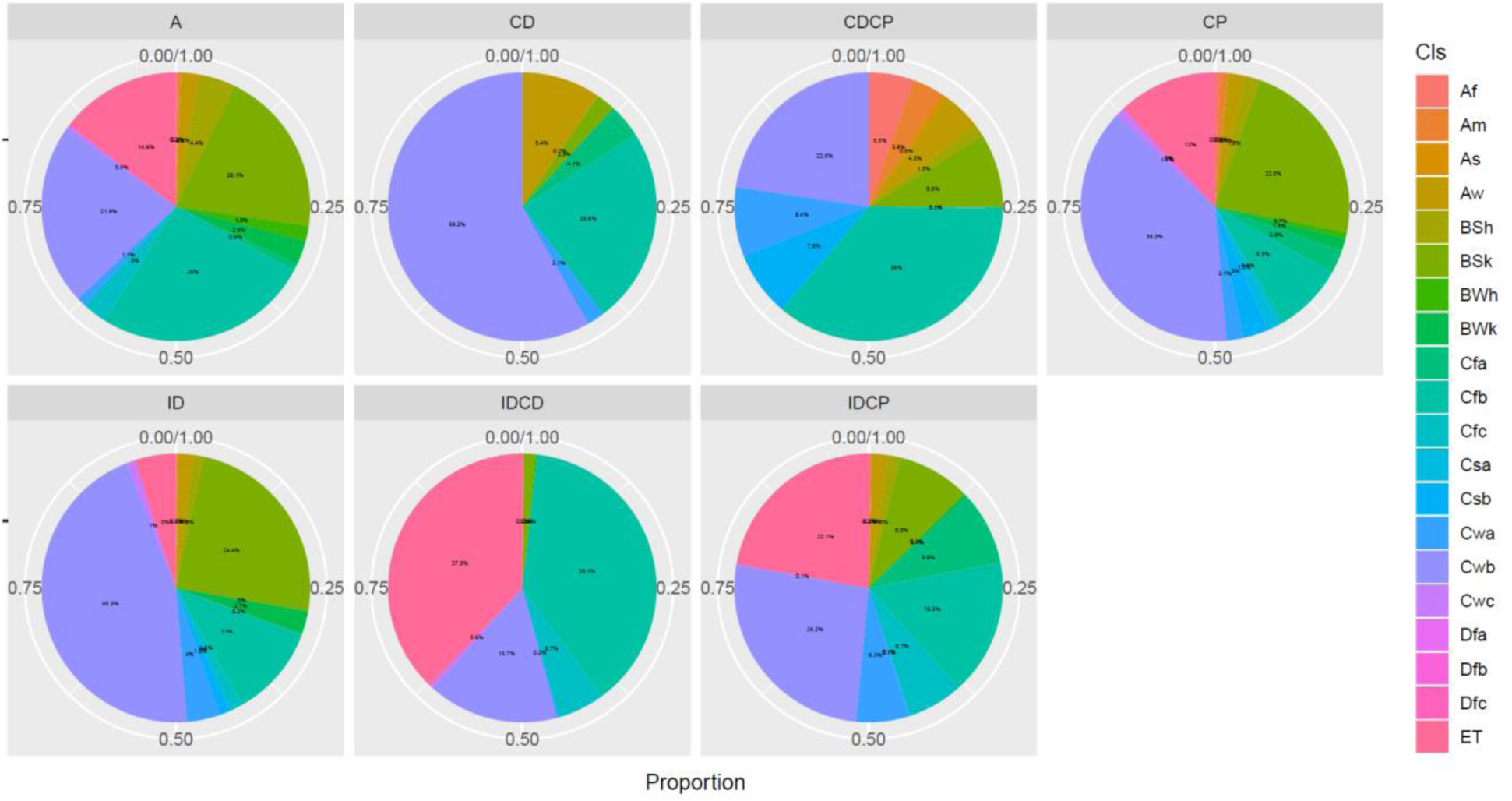
Pie chart of climate class proportions by breeding system/ploidy combinations.

